# Functional characterization of two variants in the mitochondrial topoisomerase gene TOP1MT that impact regulation of the mitochondrial genome

**DOI:** 10.1101/2021.08.02.454711

**Authors:** Iman Al Khatib, Jingti Deng, Andrew Symes, Marina Kerr, Hongliang Zhang, Sharyin Huang, Yves Pommier, Aneal Khan, Timothy E Shutt

## Abstract

*TOP1MT* encodes a mitochondrial topoisomerase that is important for mtDNA regulation, and that is involved in mitochondrial replication, transcription and translation. Two variants predicted to affect TOP1MT function (V1 - R198C and V2 - V338L) were identified by exome sequencing of a newborn with hypertrophic cardiomyopathy. As no pathogenic *TOP1MT* variants had been confirmed previously, we characterized these variants for their ability to rescue several TOP1MT functions in knockout cells. Consistent with these TOP1MT variants contributing to the patient phenotype, comprehensive characterization suggests that both variants had impaired activity. Critically, neither variant was able to restore steady state levels of mitochondrial-encoded proteins, nor reduced oxidative phosphorylation when re-expressed in TOP1MT knockout cells. However, the two variants behaved differently in some respects. While the V1 variant was better at restoring transcript levels, the V2 variant was able to restore mtDNA copy number and replication. These findings suggest that the different TOP1MT variants affect distinct TOP1MT functions. Altogether, these findings begin to provide insight into the many roles that TOP1MT plays in the maintenance and expression of the mitochondrial genome, and how impairments in this important protein may lead to human pathology.

## Introduction

Mitochondria maintain a highly reduced genome (mtDNA) that is evidence of their bacterial ancestry. In humans, the circular ∼16.5 kb genome encodes 13 proteins that are essential components of oxidative phosphorylation (OxPhos) complexes, as well as two ribosomal RNAs and 22 tRNAs required for mitochondrial translation. As such, replication, maintenance, and expression of mtDNA is important for many mitochondrial functions.

Mitochondrial genomes are present in hundreds to thousands of copies per cell and are packaged into proteinaceous structures known as nucleoids, which typically each contain one to two copies of the genome[1]. Although we do not have a complete understanding of how mtDNA abundance is maintained, mtDNA copy number is generally thought to reflect the energy production in cells, and mtDNA depletion is linked to human disease[2].

Key regulatory elements for mtDNA replication and transcription are located within an ∼1100 bp non-coding portion of the genome known as the D-loop region, due to the presence of a stable triple-stranded DNA structure that can form[3]. This D-loop comprises a short strand of DNA known as the 7S DNA that is found in a subset of mtDNA molecules and can be replicated independently of the entire genome. While the D-loop is important for mtDNA, the factors that regulate its biogenesis are poorly understood, and the exact role it plays in regulating mtDNA remains undefined. Meanwhile, expression of the mtDNA is initiated at three different promoters, a light strand promoter (LSP), and two heavy strand promoters (HSP1 and HSP2), which lead to polycistronic transcripts. Notably, LSP also plays an essential role in priming mtDNA replication[4]. Thus, the D-loop region is important to balance replication and expression of the mtDNA genome, though the exact mechanisms remain unknown. Replication, maintenance, and transcription of mtDNA is performed by proteins that are encoded in the nucleus[5-7]. The fact that mitochondria use a dedicated set of core machinery for mitochondrial transcription[6] and replication[8] highlights the fact that these processes are mechanistically distinct in mitochondria compared to the nucleus. However, there are also several proteins that function in both mitochondrial and nuclear DNA maintenance. Importantly, our understanding of how the mtDNA is regulated is still incomplete.

Topoisomerases cut and re-ligate the DNA backbone to resolve torsional strain, crossovers, and knots, and are thus important for maintenance of the circular mtDNA genome[9].

To date four of the six human topoisomerases have been found in mitochondria (*i*.*e*. TOP1MT, TOP2α, TOP2β and TOP3α). However, TOP1MT is the only topoisomerase that is found exclusively in mitochondria[10, 11], suggesting a special role for this protein in maintaining mitochondrial function. TOP1MT can relieve negative supercoiling of the mtDNA via its topoisomerase activity, which is thought to be important for its roles in mtDNA replication and transcription[12-14]. TOP1MT also binds mtDNA in the D-loop region, where it is implicated in regulating formation of the 7S DNA[12]. In addition, TOP1MT has a distinct role in mitochondrial translation[15]. Nevertheless, exactly how TOP1MT performs these various roles remains to be fully elucidated.

Here, we characterize two putative pathogenic TOP1MT variants identified in a patient who presented with hypertrophic cardiomyopathy as a newborn infant. While we show that both variants have reduced functionality, which may explain the patient pathology, they also have distinct differences that begin to shed light on the roles that TOP1MT plays in mediating mitochondrial function.

## Materials and Methods

### Consent

The methods used to identify the variants through Whole Exome Sequencing (WES) have been described previously in the MITO-FIND pipeline[16], sponsored by MitoCanada (http://mitocanada.org) and in-kind support from Discovery DNA (Calgary, Alberta, Canada). In accordance with the principles of the Declaration of Helsinki, all study participants provided informed consent. Consent was obtained following standard procedures through the University of Calgary Conjoint Research Ethics Board.

### Cell culture

HCT 116 control, KO and stably transfected cells were cultured in McCoy’s 5A (modified) media (Gibco, 16600-82) containing L-Glutamine and supplemented with 10% fetal bovine serum (FBS). Unless otherwise indicated, cells harvested for analysis were seeded at 1.5 × 10^6^ cells in 10cm plates and allowed to grow for two days, reaching approximately 75% confluence. Phoenix cells used to make retrovirus were grown in DMEM (Gibco, 11965092) supplemented with 10% FBS. All cells were maintained at 37 °C and 5% CO_2_ incubator.

### Plasmids & cloning

The pET15b/TOP1MT plasmid that has the TOP1MT open reading frame (ORF) was a gift from Dr. Yves Pommier[10]. The TOP1MT open reading frame (ORF) was cloned using InFusion cloning (Takara, Clonetech) into the pRV100G retroviral vector containing a C-terminal T2A mCherry tag, which encodes an mCherry open reading frame following self-cleaving T2A site. The TOP1MT C592T and G1012T variants were introduced using site-directed mutagenesis. An empty vector containing only mCherry was used as a control plasmid.

### Generation of stable lines generation

Stable cells expressing wild-type TOP1MT, TOP1MT variants, or empty vector controls were generated via retroviral transduction. Briefly, Phoenix cells were transfected with the retroviral plasmids, and 5 ml of the supernatant containing virus from these cells were used after 48 and 72 hrs to transduce HCT WT or TOP1MT-KO cells at 60% confluency in 100 mm plates. Following transduction, approximately 6×10^6^ cells were sorted for red fluorescence using a 130 µm nozzle on a BD FACSAria Fusion (FACSAriaIII) cytometer (BD Biosciences), supported by FACSDiva Version 8.0.1.

### PCR and sequencing

Total DNA was extracted using PureLink Genomic DNA Mini Kit (Thermo Fisher Scientific, K182001). PCR amplification of *TOP1MT* was used to confirm the success of the transduction using the forward primer CACAACAAAGGAGGTTTTCCGGAAG and the reverse primer TGCAGTCCTTCCCCAGGA. The amplified band was further sequenced using Sanger sequencing to confirm the variant status of *TOP1MT* in each line.

### Live cell imaging

For live cell imaging, 8□×□10^4^ cells were seeded on 35□mm glass bottom dishes and grown for two days. To visualize mitochondrial networks and mtDNA nucleoids, cells were simultaneously stained with MitoTracker Red (50□nM) (Thermo Fisher Scientific, M7512) and PicoGreen (Thermo Fisher Scientific, P7581) (3□μL/mL) as performed previously[17].

### Immunofluorescence staining

Cells were seeded at 1.5□×□10^4^ cells over 12□mm glass coverslips (no. 1.5) in 24 wells plates and incubated for 2□days. Subsequently, cells were fixed with 4% paraformaldehyde and stained with mitochondrial networks labeled with a primary antibody against TOMM20 (Sigma-Aldrich, HPA011562) used at 1:1000, and appropriate Alexa fluor-conjugated secondary antibodies (Thermo Fisher Scientific) at 1:1000.

### Microscopy

Fixed samples were imaged on an Olympus spinning disc confocal system (Olympus SD OSR) (UAPON 100XOTIRF/1.49 oil objective) operated by Metamorph software. A cellVivo incubation module was used to maintain cells at 37 °C and 5% CO2 during live cell imaging.

### Image analysis for mtDNA nucleoids

Nucleoid counts and size measurements were obtained using the particle analysis tool in ImageJ FIJI after binarizing the images[18]. Surface area (size) of each nucleoid and the total nucleoid count per cell were automatically determined for a manually selected region of interest containing the entire mitochondrial network, but excluding the nucleus. The analyses were performed on at least 10 cells for each of the indicated cell lines. For live cells stained with PicoGreen, a violin plot was used to represent the distribution of mtDNA nucleoid sizes and counts for the indicated cell lines. For fixed cells stained with anti-DNA antibody, nucleoid sizes are presented as the average size ± SEM. The non-parametric Kolmogorov-Smirnov (K-S) test was used to determine statistical significance regarding the distribution of counted nucleoids.

### Image analysis for mitochondrial networks

Mitochondrial network morphology was qualitatively analyzed by classifying networks into three categories, fragmented (predominantly small mitochondrial fragments), intermediate (cells containing a mixture of short fragments and elongated networks) and fused (elongated, interconnected networks with few to no short fragments)[17]. For each cell line investigated, at least 50 cells were scored. Morphology analyses were performed on 3 independent replicates. The results shown represent mean□±□SEM, and *P* values were based on unpaired, 2-tailed Student’s *t*-tests. Average mitochondrial length was measured using ImageJ FIJI. Briefly, background signal was subtracted, images of mitochondrial networks were skeletonized, and the mitochondrial length was obtained using the analyze skeleton function in ImageJ FIJI. The sum of the lengths of all branches of a mitochondrion was evaluated as the total length of that mitochondrion in every image[19]. For each of WT and KO cells, at least 20 cells were evaluated. The results shown are one of three independent biological replicates with the same trends showing mean□±□SEM, and p values based on unpaired, 2-tailed Student’s t-tests.

### Long-range PCR

To examine mtDNA deletions, we amplified a 16.2 kb fragment representing nearly the full length mtDNA, as described previously [20] using the following primers (1482–1516 F: ACCGCCCGTCACCCTCCTCAAGTATACTTCAAAGG; 1180–1146 R: ACCGCCAGGTCCTTTGAGTTTTAAGCTGTGGCTCG). Long range PCR reactions were performed using the Takara LA Taq polymerase (Takara Bio, RR002M), with 250 ng genomic DNA, 200 nM forward and reverse primers. The PCR cycling conditions were as follows: 94 °C for 1 min; 98 °C for 10 s and 68 °C for 11 min (30 cycles); and a final extension cycle at 72 °C for 10 min. PCR products were visualized by electrophoresis on a 0.8% agarose gel, run for approximately 16 h at 20 V.

### mtDNA copy number analysis

Total DNA was isolated from cultured cells using the PureLink Genomic DNA Mini Kit (Thermo Fisher Scientific, K182001) according to the manufacturer’s protocol, and 50-100 ng of DNA was used for quantitative PCR (QPCR) to determine mtDNA copy number, as reported previously[21]. Primers used are shown in Table 1. Briefly, mtDNA or 18S DNA were amplified using PowerUp SYBR Green Master Mix (Thermo Fisher Scientific, A25742) in the QuantStudio 6 Flex Real-Time PCR system (Thermo Fisher Scientific) machine, and the delta delta Ct method was used to determine mtDNA copy number relative to the 18S. Reactions were performed in triplicate and mtDNA copy number analysis was performed on at least three independent biological replicates. Data is presented as mean□±□SEM and unpaired, 2-tailed Student’s *t*-tests were used to determine statistical significance.

**Table 1:**
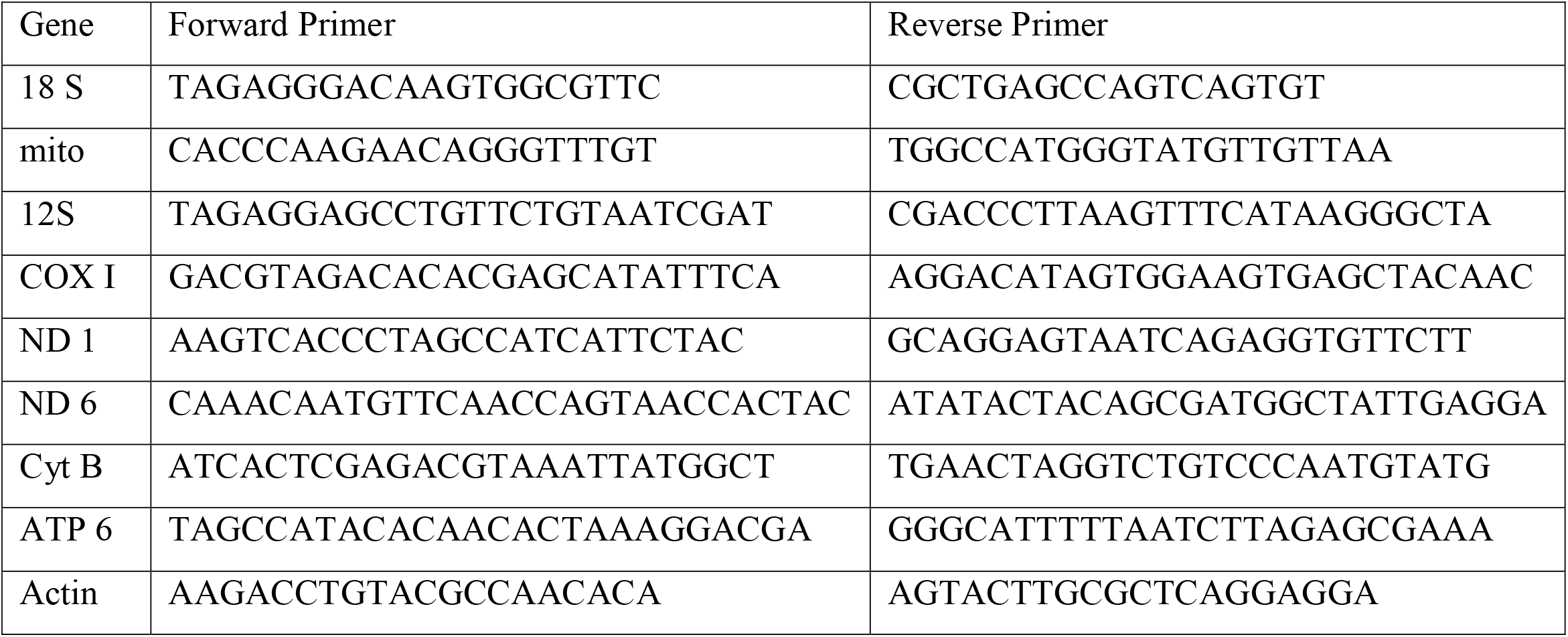
Primers used for quantitative PCR for detection of different genes as indicated where 18S and mito were used for detection of mtDNA copy number and the rest for RNA expression of various genes

### Ethidium bromide depletion/repletion

Cells were treated with 1µg/ml Ethidium bromide (Fisher Scientific, 15-585-011) for three days, as described previously[22], at which point fresh media was added. Cell pellets were collected at days 0, 3, 6 and 8, for DNA purification and copy number analysis.

### Western

Cell pellets were collected after trypsinization, washed with 1× phosphate buffered saline (PBS) and lysed with RIPA buffer (Thermo Scientific™, 89900) containing protease inhibitors. 20-50 µg of total cell lysates were resolved on SDS-PAGE gels and transferred onto PVDF membranes. Blots were probed with the following antibodies at the indicated dilutions; OxPhos antibody cocktail (Abcam, ab110411; 1:500 dilution), anti-Actin (Sigma, A5316; 1:1000), anti-NDUFB6 (Abcam, ab110244; 1:1000), anti-MTCO2 (Abcam, ab110258; 1:1000), anti-GAPDH (Millipore Sigma, ABS16; 1:1000), anti-VDAC1 (Abcam, ab14734; 1:1000), anti-TOP1MT-3 (Developmental Studies Hybridoma Bank, CPTC-TOP1MT-3; 1:200). Appropriate horseradish peroxidase (HRP)-conjugated secondary antibodies were used at 1:3000 as follows: goat anti rabbit IgG, HRP linked Antibody (Cell Signaling Technology, 7074S), or goat anti-mouse IgG-HRP (Santa Cruz Biotechnology, sc-2055). Blots were incubated with Clarity ECL substrate (Biorad, 1705061) and imaged using an Amersham Imager AI600.

### Southwestern blot

To block nuclear DNA replication, cells were treated for 4 hours with 10 µM Aphidicolin (HelloBio Ltd, Bristol UK). Next 50 uM BrdU (Invitrogen, fisher scientific) was added for 24 hours, and total DNA isolated from 2 × 10^7^ cells using the E.Z.N.A Tissue DNA kit. To help identify supercoiled, relaxed or linear mtDNA, 1µg of total DNA was treated with 10 U of Topoisomerase I (New England Biolabs) for 1 hour at 37°C, or 20 U of *BamHI* (New England Biolabs) for 1 hour at 37°C. DNA samples (1ug) were then separated on a 0.45% agarose gel (Biotechnology VWR, life science) at 30V 18-20 hours at 4°C. The gel was washed with water prior to soaking for 45-60 mins in denaturing buffer (0.5N NaOH, 1.5M NaCl). After another wash in water, the gel was soaked for 30 mins in 10X saline-sodium citrate (SSC). DNA was then transferred to a PVDF membrane (Bio-Rad, Cat# 1620177) using capillary action as described previously (Nature protocols, 518, 2006). The PVDF membrane was washed in 6X SSC for two minutes, and DNA immobilized on the membrane by UV-crosslinking (UV Stratalinker 1800). The membrane was blocked with 10% milk for 1h, prior to 2 h incubation for at room temperature with anti-BrdU antibody (Becton Dickinson Immunocytometry Systems) at 1:2000 dilution in TBST, followed by incubation with horseradish peroxidase (HRP)-conjugated goat polyclonal anti-mouse IgG (1:5,000 dilution, EMD Millipore Etobicoke, Ontario, Canada). Supersignal West Femto enhanced chemiluminescence substrate (Thermo Scientific) was used and blots imaged using the chemiluminescence imaging analyzer (LAS3000mini; Fujifilm, Tokyo, Japan).

### Mitochondrial respiration

Mitochondrial bioenergetic profiles were obtained using a Seahorse XFe24 Extracellular Flux Analyzer (Agilent Technologies, Inc) with the Seahorse XF Cell Mito Stress Test. Briefly, 8 ×□10^4^ cells/well were seeded in an XF24 microplate and incubated at 37□°C, 5% CO_2_ for 48□h. Subsequently, growth media was replaced with assay media supplemented with D-Glucose (25□mM), sodium pyruvate (2□mM) and L-Glutamine (4□mM). Oxygen consumption rates were measured following sequential injection of the following compounds into each well: oligomycin (1□μg/mL) (Enzo Life Sciences, BML-CM111), carbonyl cyanide 4-(trifluoromethoxy) phenylhydrazone (FCCP, 1□μM) (Enzo life Sciences, BML-CM120) and Antimycin A (1□μM) (Sigma Aldrich, A8674). Upon completion of the assay, protein concentrations from each well were measured by BCA assay (Thermo Fisher Scientific, 23225) and used to normalize data.

### RNA extraction and quantification

RNA was extracted from cell pellets using the E.Z.N.A.® HP Total RNA Kit, Omega Bio-tek® (VWR Scientific, CA101414-850), according to the manufacturer’s protocol.

Following extraction, cDNA was generated from 7.5 µg of RNA using the iScript™ Advanced cDNA Synthesis Kit (Biorad, 1725038), and 15-20 ng of cDNA was then used for quantitative PCR with PowerUp SYBR Green Master Mix (Thermo Fisher Scientific, A25742) in the QuantStudio 6 Flex Real-Time PCR system (Thermo Fisher Scientific) machine. The delta delta Ct method was used to determine the relative expression of various transcripts using the primers indicated in Table 1 relative to Actin expression. Reactions were performed in triplicate and mtDNA copy number analysis was performed on at least three independent biological replicates. Data is presented as mean□±□SEM and unpaired, 2-tailed Student’s *t*-tests were used to determine statistical significance.

### Structural modeling

Structural models for the TOP1MT variants were generated via RoseTTAFold, a deep learning solution to predicting protein structures, which was conducted on the Robetta web server(robetta.bakerlab.org)[23].

## Results

### Patient Description

We report a newborn male infant who at birth presented with hypertrophic cardiomyopathy characterized primarily by concentric left ventricular hypertrophy with biventricular hypertrophy and prominent trabeculations. The patient was one of two non-identical dichorionic/diamniotic twins born to a 50-year-old mother. Both parents were of Nigerian ancestry. Whereas the other twin was healthy, the patient was born with severe hypertrophic obstructive cardiomyopathy. While the mother had a history of gestational diabetes, the cardiac hypertrophy in the patient was severe in nature, and being present in only one of the twins, was considered needing further investigations, and prompted testing through whole exome sequencing (WES). Additional clinical workup of the patient at birth showed hyperglycaemia, which could be due to stress of the unusual birth, and hepatomegaly. The acylcarnitine profile showed elevated C18:2, C16 and C18 acyl-carnitine species. However, WES did not identify any variants in genes associated with trifunctional protein deficiency or beta-oxidation that would be considered biologically impactful. There was no elevation in blood lactate or alanine. The cardiac hypertrophy responded to treatment with a beta-blocker, propranolol, and the infant was discharged. The family moved to a different country, and no follow up scans or other samples from the family were available. At a 6-month follow-up contact, the baby was reported to be healthy, but had not seen a cardiologist or had further testing in his area of residence.

### Variant Identification

Whole exome sequencing revealed the patient had compound heterozygous variants in *TOP1MT* (NM_052963.2). The first variant, V1, was found at position C592T, and causes an arginine to leucine change at amino acid 198 (Arg198Cys). The second variant, V2, was found at G1012T, and yields a valine to leucine change at residue 338 (Val338Leu). Parental DNA was not available for segregation testing. Both identified variants occur at highly conserved sites in the core domain of TOP1MT (Fig 1a) and were predicted to be deleterious and damaging with allele frequencies of 0.000527 for V1 and 0.00987 for V2 in Genome Aggregation Database (GnomAD). Notably, the V2 variant was reported previously as a potentially deleterious genomic missense variant for TOP1MT[24]. Nonetheless, as *TOP1MT* is not formally recognized as a human disease gene, additional functional characterization of these TOP1MT variants was required to investigate their potential pathogenicity.

**Figure 1.**
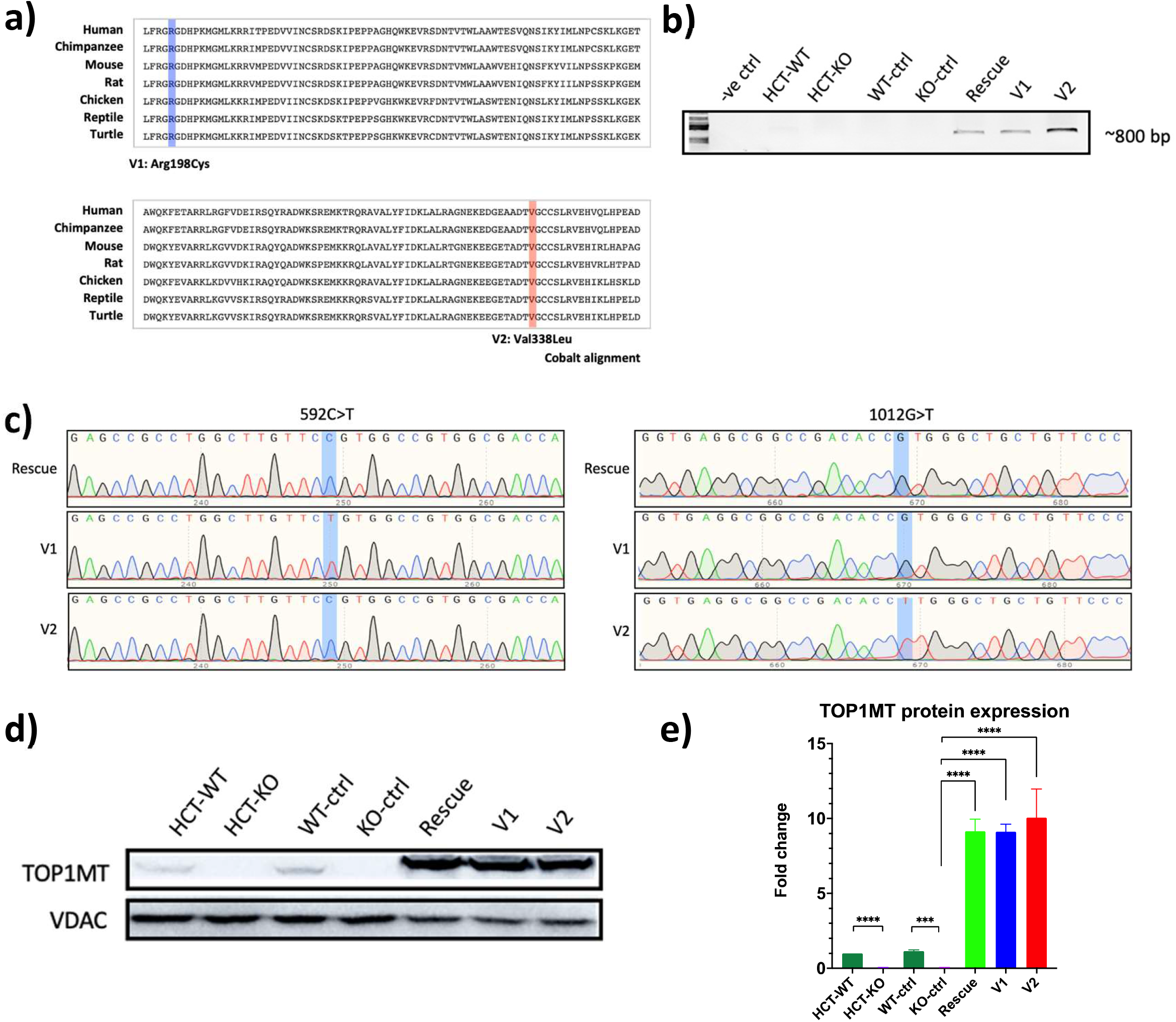
Generation of stable lines expressing wild-type (WT) or mutant forms of TOP1MT, as well as control lines for both WT and KO cells. **a)** Amino acid alignments of TOP1MT proteins from a variety of vertebrate species generated using COBALT shows the conservation of the amino-acids variants identified in the patient. **b)** PCR amplification targeting the coding region of the viral-encoded *TOP1MT* open reading frame, spanning patient variants, confirms amplicons are present only in HCT-KO cell lines stably re-expressing *TOP1MT*, namely the Rescue, V1 and V2 lines. **c)** Chromatograms from Sanger sequencing of the amplicons produced from (b), for the 592C and 1012G residues (shaded), confirms the identity of the cell lines. **d)** Western blots probed for TOP1MT antibody confirms the HCT-KO line lacks TOP1MT and reveals TOP1MT expression in stably transduced cell lines. VDAC was probed as a load control. **e)** Quantification of western blots from 4 replicates show the fold change in protein abundance compared to VDAC control from four different replicates. All statistical analysis were done using unpaired student t-test; where error bars represent standard error of mean and p values ***<0.001 and ****<0.0001.

### Cell Line Generation

To examine the functionality of the V1 and V2 variants, we characterized the ability of these variants to rescue functions when re-expressed in an HCT116 cell line where *TOP1MT* was knocked via CRISPR-Cas9[22]. Starting from HCT116 wild-type or HCT116 *TOP1MT* KO cells (labelled HCT-WT or HCT-KO, respectively), we generated five different stable cell lines using retroviral constructs for TOP1MT-P2A-mCherry, or an empty vector control. As controls to eliminate the effect of retroviral transduction and mCherry expression, the HCT-WT and HCT-KO cells were transduced with the T2A-mCherry empty vector (noted as WT-ctrl and KO-ctrl). Importantly, the empty vector WT-ctrl and KO-ctrl lines behaved similar to their untransduced parental HCT-WT or HCT-KO lines with respect to the TOP1MT functions investigated throughout this paper, indicating that the viral transduction itself had no major impact. Next, to examine the rescue effects of re-expressing wild-type TOP1MT in KO cells, a third cell line was generated by re-expressing the wild-type form of TOP1MT (indicated as Rescue). Finally, to evaluate the ability of the patient variants to rescue loss of TOP1MT function in KO cells, two additional cell lines expressing either the V1 or V2 variants were established (referred to as V1 or V2, respectively).

Viral transduction of cells was confirmed by visualizing mCherry expression via confocal microscopy and cells expressing mCherry were sorted by flow cytometry to select the population of transduced cells. The transduction of WT, V1 and V2 TOP1MT was further confirmed by PCR amplification of the TOP1MT open reading frame (Fig 1b), and the variant status of the Rescue, V1 and V2 lines was confirmed by DNA sequencing (Fig 1c).

Next, we confirmed the expression of TOP1MT proteins by immunoblotting (Fig 1d & 1e). A band of the predicted size of ∼70 kDa corresponding to TOP1MT protein was observed in HCT-WT and WT-ctrl cells, which is missing from the HCT-KO and KO-ctrl cells. A band migrating slightly slower was observed in the Rescue, V1, and V2 lines (Fig 1d), reflecting extra sequence from the T2A site.

Immunofluorescence imaging confirmed the protein expression findings from the western blotting, and showed that WT, V1, and V2 TOP1MT proteins are indeed localized to mitochondria (Fig 2). Having established these cell lines, we then looked to determine whether phenotypic differences in HCT-KO and HCT-WT cells without viral transduction (Fig S1) could be rescued by re-expression of WT, V1, or V2 TOP1MT variants. Thus, we examined the several cellular parameters.

**Figure 2.**
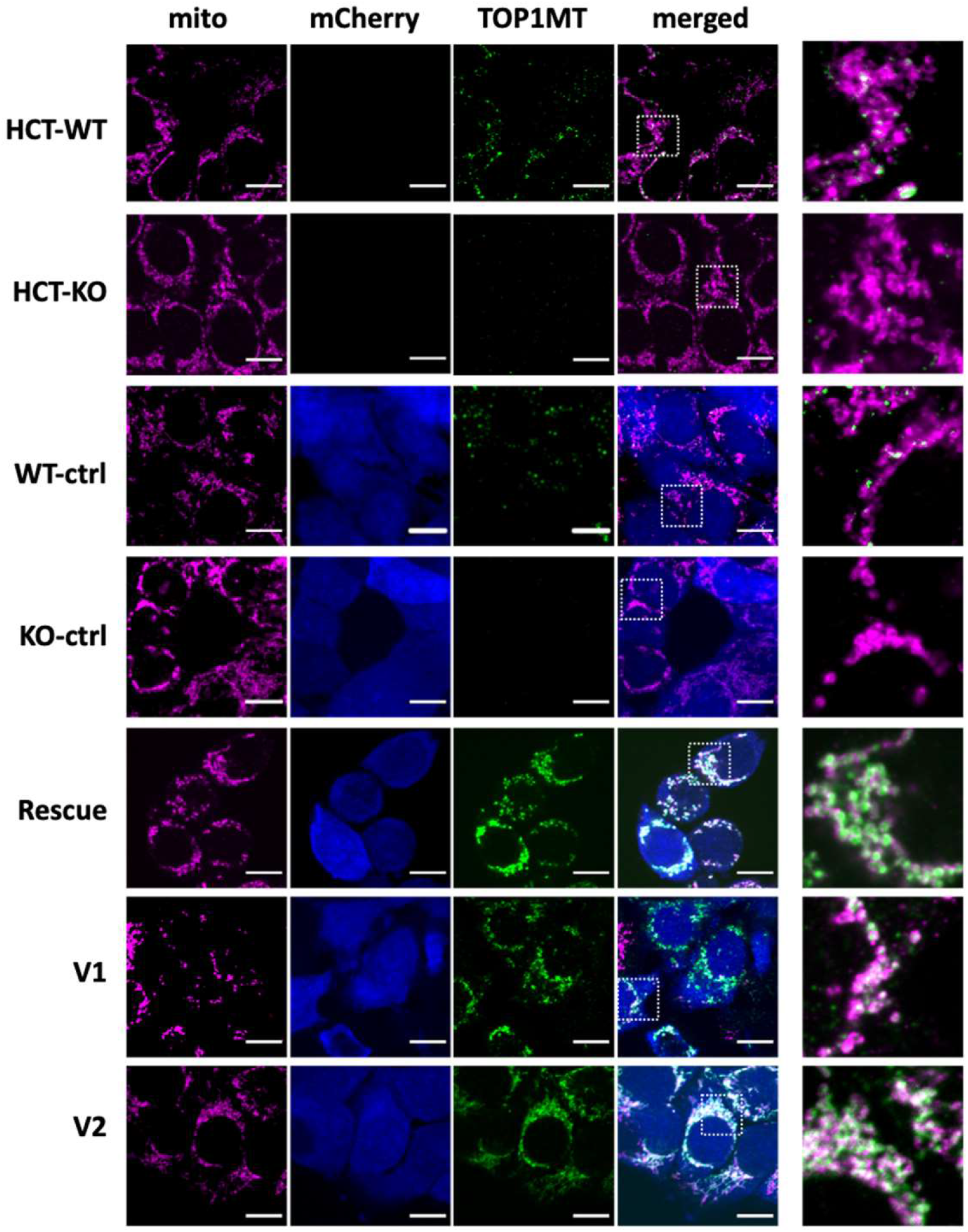
Representative confocal images of TOP1MT localization in stable lines. Fixed Cells were stained for immunofluorescence with anti-TOMM20 (mitochondria, magenta) and anti-TOP1MT (green) primary antibodies. Viral transduction was confirmed by expression of mCherry (Blue). The final column shows the zoomed-inn region indicated in the merged column. Images confirm lack of TOP1MT in HCT-KO cells and show the mitochondrial localization of TOP1MT-WT and both V1 and V2 patient variants. Scale bar represent 10 μm.

### Nucleoids

Given the role of TOP1MT as a mitochondrial DNA topoisomerase that reverses negative supercoiling of mtDNA[25], we examined the effect of TOP1MT loss on nucleoids. We employed confocal microscopy immunofluorescence to visualize nucleoids in live cells labelled with PicoGreen (Fig 3a). Although there was a trend towards an increase in the average number of nucleoids per cell in the KO cells, this was not statistically significant (Fig 3b). However, KO cells did show a significant ∼30% reduction in nucleoid size, consistent with increased negative supercoiling (Fig 3c). The change in nucleoid size was not due to the accumulation of large mtDNA deletions, as long-range PCR confirmed that there were no mtDNA deletions in either the HCT ctrl or HCT-KO cell lines (Fig 3d). Importantly, a control cell line bearing deletions was used as a positive control to show the assay can detect mtDNA deletions.

**Figure 3.**
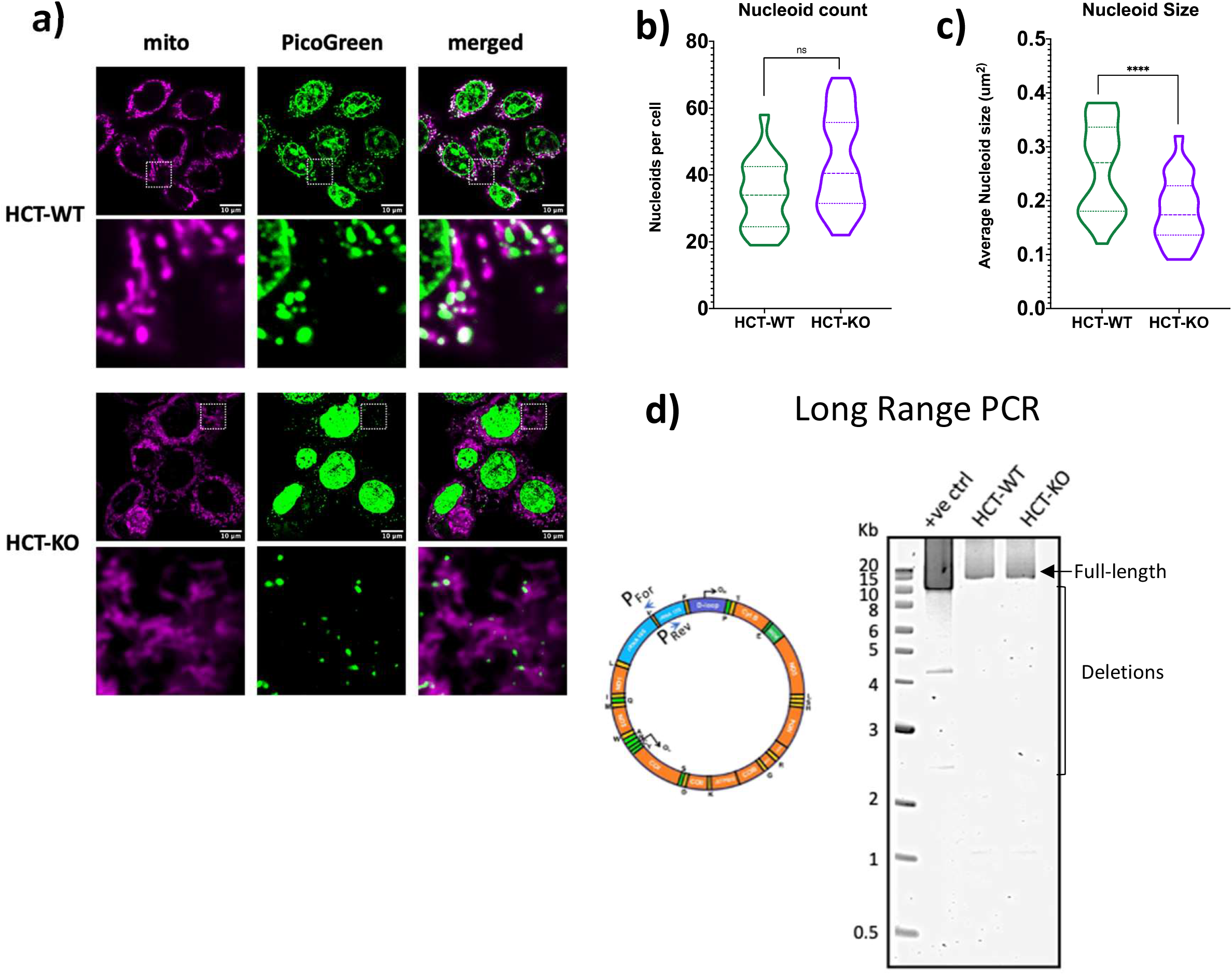
Analysis of mitochondrial nucleoids and mtDNA deletions in control and TOP1MT-KO cells. **a)** Representative confocal images of live cells stained with MitoTracker Far Red (mitochondria) and PicoGreen (dsDNA: nuclear and mtDNA) where the final column shows the merged image between mitochondria and dsDNA. Scalebars represent 10μm. A zoomed-in region is shown on the lower row corresponding to the represented image above for both HCT-WT and HCT-KO cells. Violin plots showing quantification of mtDNA nucleoids from 25 cells stained as in (a), represented as nucleoid count per cell **(b)** and average nucleoid size per cell **(c)** P-values were determined by a Kolmogorov-Smirnov test for all measured nucleoids, where ‘ns’ signifies no significant differences between indicated groups and ****<0.0001. **d)** Long range PCR of mtDNA in HCT-WT and KO cells show only the full-length 16.3 kb product, indicating no mtDNA deletions are detectable. DNA form a cell harbouring a mtDNA deletion was used as a +ve control to show the protocol detects deletions.

We next asked whether the reduced nucleoid size in the HCT-KO cells could be rescued by re-expressing TOP1MT. Critically, overexpression of the WT TOP1MT protein reversed the nucleoid size phenotype, whereas the overexpression of V1 and V2 variants led to intermediate nucleoid sizes, with V1 showing better rescue compared to V2 (Fig 4a & 4b). Since the ability of PicoGreen dye to bind mtDNA can be affected by supercoiling[26-28], which may have skewed our results, we also examined nucleoid sizes in fixed cells labelled with anti-DNA antibody, and similar results were obtained (Fig S2).

**Figure 4.**
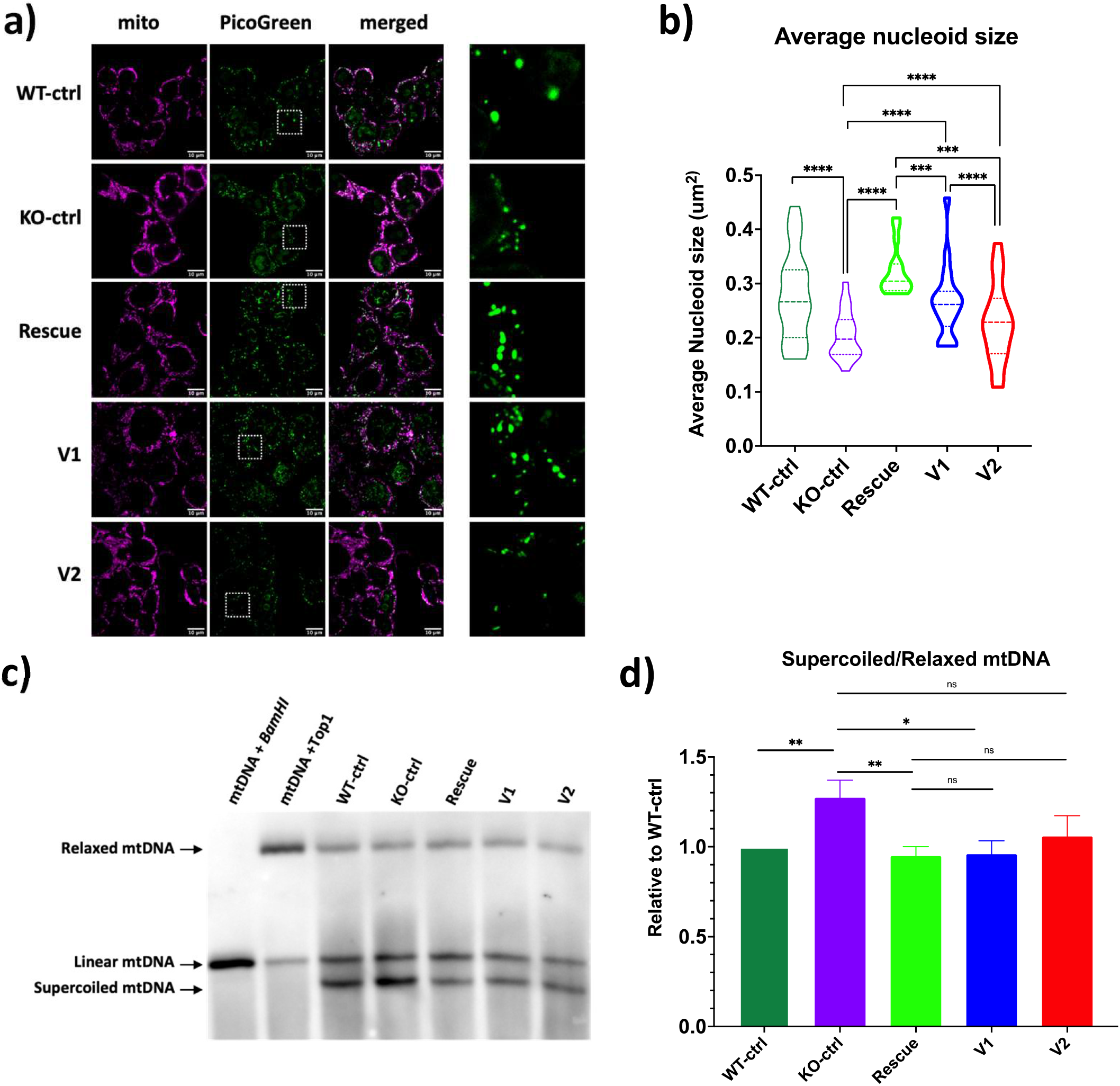
Nucleoid sizes and mtDNA supercoiling in TOP1MT cell lines. **a)** Representative confocal images stained of live cells with MitoTracker Far Red (mitochondria) and PicoGreen (dsDNA: mtDNA nucleoids and nucleus). Merged column represents the combined image of mitochondria (purple) and dsDNA (green), where the final column shows the zoomed-in region of nucleoids for size comparison. Scalebars represent 10 μm. **b)** Quantification of mtDNA nucleoid sizes in cells stained as in (a). Data represents average nucleoid sizes from at least 15 cells for each group. P-values were determined by a Kolmogorov-Smirnov test for all measured nucleoids, with ***<0.001 and ****<0.0001. **c)** Representative southwestern blots shows increased levels of the supercoiled mtDNA band in KO ctrl cells and intermediate rescue by V1 and V2. Control samples treated with *BamHI* or Topoisomerase I (Top) identify relaxed, supercoiled and linear mtDNA species. **d)** Quantification of southwestern blots of different mtDNA forms as in (a), from four independent experiments, as a ratio of supercoiled to relaxed mtDNA. All statistical analysis were done using unpaired student t-test and p values *<0.05, **<0.01 and ‘ns’ signifies no significant differences between indicated groups. Error bars represent standard error of mean.

While the decreased nucleoid size in KO cells is consistent with increased supercoiling, changes in nucleoid size could also be due to alterations in clumping or aggregation of multiple nucleoids, which cannot be visualized with standard confocal microscopy. Thus, we also utilized a southwestern blot approach to look at the mtDNA topology. In this approach, BrdU-labelled mtDNA is separated on an agarose gel based on its conformation, transferred to a membrane, and detected by an anti-BrdU antibody [29, 30]. Notably, nuclear DNA labelling is eliminated by the use of Aphidicolin, an inhibitor of DNA polymerase[31]. Bands representing, relaxed circular mtDNA, linear mtDNA, and supercoiled circular mtDNA [32], were confirmed by control samples treated with *BamHI*, which cuts mtDNA once creating a linear molecule, or *E. coli* topoisomerase, which relaxes supercoils. When we quantified the ratio of supercoiled to relaxed mtDNA, we confirmed that cells lacking TOP1MT had elevated mtDNA supercoiling, which was reversed upon re-expressing wild-type TOP1MT (Fig 4c & 4d). Similar to the results obtained in nucleoid size analysis, the increased supercoiling was rescued by both V1 than V2 variants.

### mtDNA copy number and replication

Although the average number of nucleoids/cell quantified by imaging was not significantly different (Fig 3b), previous reports have shown changes in mtDNA copy number under certain conditions of TOP1MT impairment[22]. Thus, we employed quantitative PCR as a more sensitive assay to look at mtDNA copy number. We found that mtDNA copy number was significantly elevated by ∼40% in KO-ctrl cells compared to WT-ctrl cells (Fig 5a), and that this was corrected in the Rescue cells re-expressing WT TOP1MT. Moreover, while expression of V1 did not restore the copy number, expression of V2 did provide rescue.

**Figure 5.**
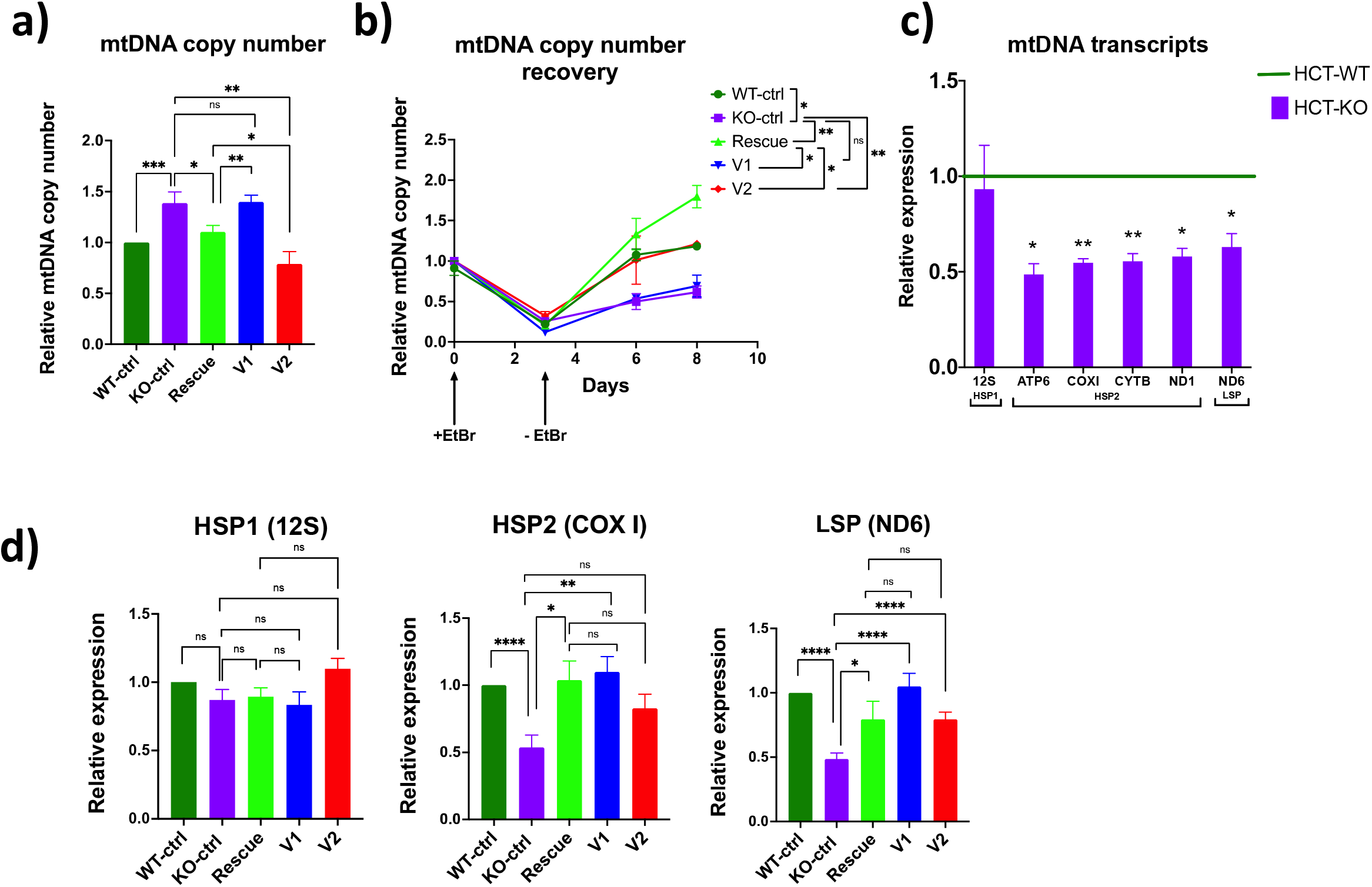
mtDNA replication and expression in TOP1MT rescue cell lines. **a)** Steady state copy mtDNA number, normalized to 18S, obtained via quantitative PCR. **b)** Recovery of mtDNA copy number, as determined in a), following three days depletion with EtBr. **c)** Quantification of mtDNA transcripts in HCT-WT and HCT-KO cell lines via quantitative RT-PCR for the indicated genes from the indicated mtDNA promoters: HSP1, LSP and HSP2. **(d)** Quantification of mtDNA transcripts in stable TOP1mt rescue lines for HSP1 (12S), HSP2 (COX I) and LSP (ND6). Error bars represent standard error of mean. P-values were determined by unpaired student t-test such that *<0.05, **<0.01, ***<0.001 and ****<0.0001. ‘ns’ signifies no significant differences between indicated groups. Data represents at least three independent biological replicates.

In addition to examining steady state mtDNA levels, we also examined the ability of the mtDNA to replicate. A common assay for mtDNA replication is a depletion/repletion assay, whereby the intercalating agent ethidium bromide blocks mtDNA replication leading to copy number depletion[33, 34]. Subsequent removal of ethidium bromide allows mtDNA replication such that repletion of mtDNA copy number can be measured[35]. Similar to previous reports[22, 36], we found that mtDNA repletion in KO-ctrl cells is significantly reduced compared to WT-ctrl (Fig 5b). Meanwhile, although re-expression of WT TOP1MT increased repletion, above endogenous levels of the WT cells, the V1 and V2 variants had different effects. The V1 variant showed no increase in repletion, while the V2 variant showed improved repletion, similar to WT-ctrl cells, but still reduced compared to the Rescue cells.

### Mitochondrial Transcription

As transcription of the mtDNA genome is intricately linked to mtDNA replication and is also influenced by mtDNA supercoiling mediated by TOP1MT[13, 14], we examined the steady-state levels of mtDNA transcripts. Unexpectedly, despite the finding that mtDNA copy number was higher in non-transduced HCT-KO cells, mtDNA transcripts from LSP and HSP2 promoters were significantly lower in these cells compared to non-transduced HCT-WT (Fig 5c). Meanwhile, levels of the rRNA transcripts from HSP1 were not affected.

As the reduced transcript levels that we observed contradicted previous reports showing increased levels of mtDNA-encoded transcripts in cells lacking TOP1MT[14, 25, 37, 38], we examined the relative transcript levels in cells grown at different seeding densities (Fig S3). We found that at lower seeding densities, similar to the experimental conditions used in this paper, HCT-KO cells had reduced levels of mtDNA-encoded transcripts. Conversely, at higher seeding densities where cells were confluent upon harvesting for experimentation, HCT-KO cells had increased levels of mtDNA-encoded transcripts.

Importantly, transcript levels for HSP2 and LSP were restored to normal by re-expressing WT TOP1MT in the Rescue cells (Fig 5d). With respect to the effect of the pathogenic variants, re-expression in the V1 cells was able to rescue both HSP2 and LSP transcript levels. Meanwhile, the V2 cells maintained lower HSP2 transcript levels but did show some rescue for LSP.

### Expression of Mitochondrial Proteins

As TOP1MT can also influence mitochondrial translation, we investigated the expression of OxPhos proteins in our cell lines. We found several intriguing differences when we examined the expression of different OxPhos complex subunits (Fig 6a). Most notably, expression of Complex I subunits NDUFB8 (Fig 6b) and ND6 (Fig 6g), Complex II subunit SDHB (Fig 6c), Complex III subunit UQCRC2 (Fig 6d), complex IV subunit COXII (Fig 6e) and Complex V subunit ATP5A (Fig 6f) showed similar trends with reduced expression in KO-ctrl cells compared to WT-ctrl cells. Although we did not see a significant decrease in UQCRC2 (Fig 6d) in these experiments, we did observe a decrease in the parent HCT-KO compared to HCT-WT cells (Fig S1). Importantly the expression of all these proteins in the Rescue cell line was restored, sometimes even increased above WT-ctrl levels. Meanwhile, the expression of these proteins in the V1 and V2 lines showed variable amounts of rescue. Generally, expression of OxPhos proteins in the V1 and V2 lines tended to be less than that in the Rescue line. Critically, there was little to no rescue for the mtDNA encoded proteins ND6 and COXII, indicating that these variants could not restore mitochondrial translation.

**Figure 6.**
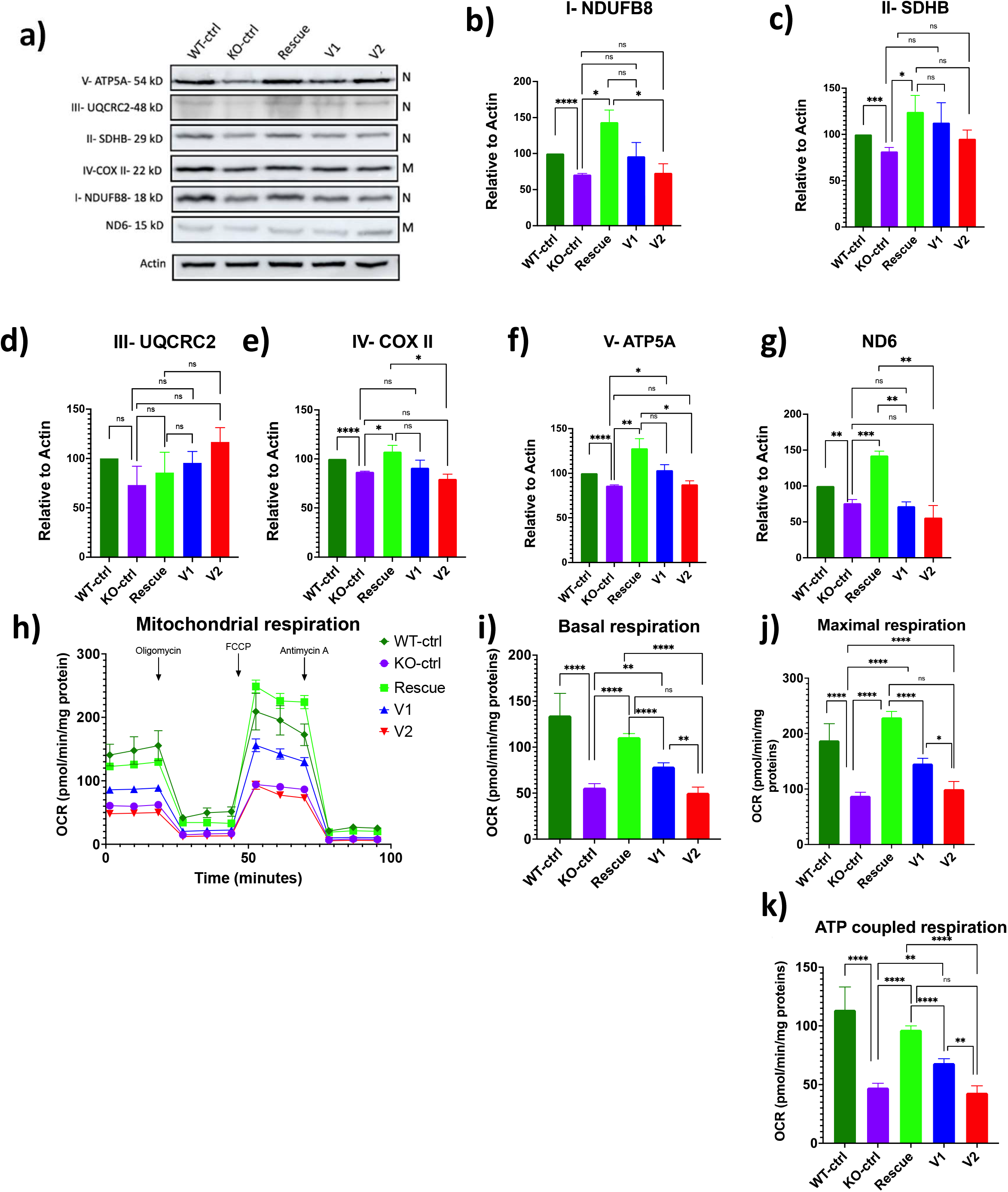
Mitochondrial protein expression and oxygen consumption in TOP1MT rescue lines. **a)** Representative western blots of the following nuclear encoded (N) or mitochondrial encoded (M) OxPhos complex proteins: NDUFB8 (Complex I-N), SDHB (Complex II-N), UQCRC2 (Complex III-N), COXII (Complex IV-M), ATP5A (Complex V-N) and ND6 (Complex I-M). **b-g)** Quantification of indicated proteins from at least three independent biological replicates of protein levels determined by western blot as in a). Expression was normalized to expression the Actin load control. **h)** Oxygen consumption rates (OCR) over time analyzed in the indicated cell lines using the Seahorse XF24 extracellular flux analyzer. The OCR data was used to calculate basal respiration **(i)**, maximal respiration **(j)** and ATP production **(k)**. Data presented is from two independent biological replicates, each with 5 technical replicates. Error bars represent standard error of mean. P-values were determined by an unpaired student t-test with *<0.05, **<0.01, ***<0.001 and ****<0.0001. ‘ns’ signifies no significant differences between indicated groups.

### Mitochondrial Respiration

Given these changes in levels of OxPhos complex protein expression, we next investigated mitochondrial respiration in our cells (Fig 6h). Similar to previous work[15], we found that both basal respiration (Fig 6i), maximal respiration (Fig 6j), and ATP-coupled respiration (Fig 6k), were all reduced in KO-ctrl cells. While re-expression of WT TOP1MT in the Rescue cells showed restoration of all these parameters, the patient variants had different effects. Expression of V1 provided partial rescue, while the V2 variant provided no rescue at all.

### Mitochondrial Morphology

Mitochondrial morphology can change in response to different stresses and is also important for the regulation of mtDNA. Thus, as another measure of mitochondrial fitness, we decided to look at mitochondrial morphology in our stable cell lines using confocal microscopy (Fig 7a). We observed a marked increase in the number of cells with hyperfused mitochondrial networks in HCT-KO cells compared to HCT-WT cells (Fig 7b & 7c). We also found that expression of wild-type TOP1MT restored mitochondrial morphology in the Rescue cells, whereas the patient variants had an intermediate effect on mitochondrial morphology in the V1 and V2 cell lines (Fig 7d & 7e).

**Figure 7.**
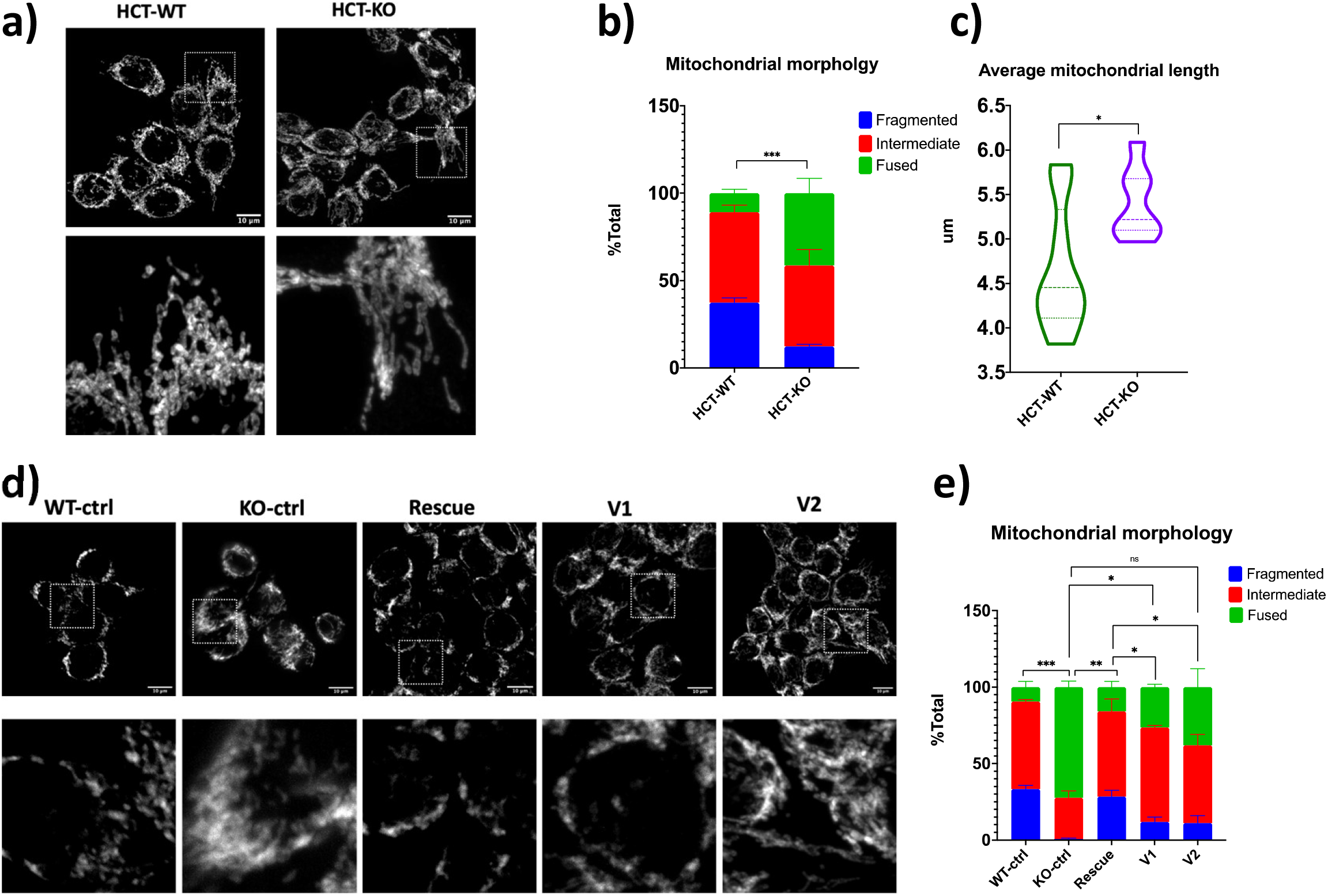
Mitochondrial network morphology in TOP1MT lines. **a**,**d)** Representative confocal images of mitochondrial networks from fixed cells and stained for immunofluorescence with the outer membrane protein TOMM20. Scalebars are 10 μm. **b**,**e)** Qualitative quantification of mitochondrial morphology for cells stained as in (a) and (e), respectively. Analysis for each line was performed on three technical replicates of at least 50 cells each. **c)** Quantitative analysis of average mitochondrial length from 20 cells as in (a), determined using ImageJ, confirms the qualitative analysis in (b). Error bars represent standard error of mean. P-values for the percentage of cells with fused mitochondrial networks (b and e), or average mitochondrial length (c), were determined by an unpaired student t-test with *<0.05, **<0.01, ***<0.001 and ‘ns’ signifies no significant differences between indicated groups.

### Structural modeling of the TOP1MT variants

In an effort to understand how the V1 and V2 variants impact the various functions of TOP1MT, we performed structural modelling of the protein variants. Our modelling generated by RoseTTAFold[23] shows that TOP1MT is predicted to bind dsDNA tightly through a barrel-like structure in the core domain of the protein (Fig 8). In this case we used an mtDNA sequence that was suggested previously to bind TOP1MT[12, 39]. Intriguingly, both the R198 and V338 residues are located within the putative DNA binding socket.

**Figure 8.**
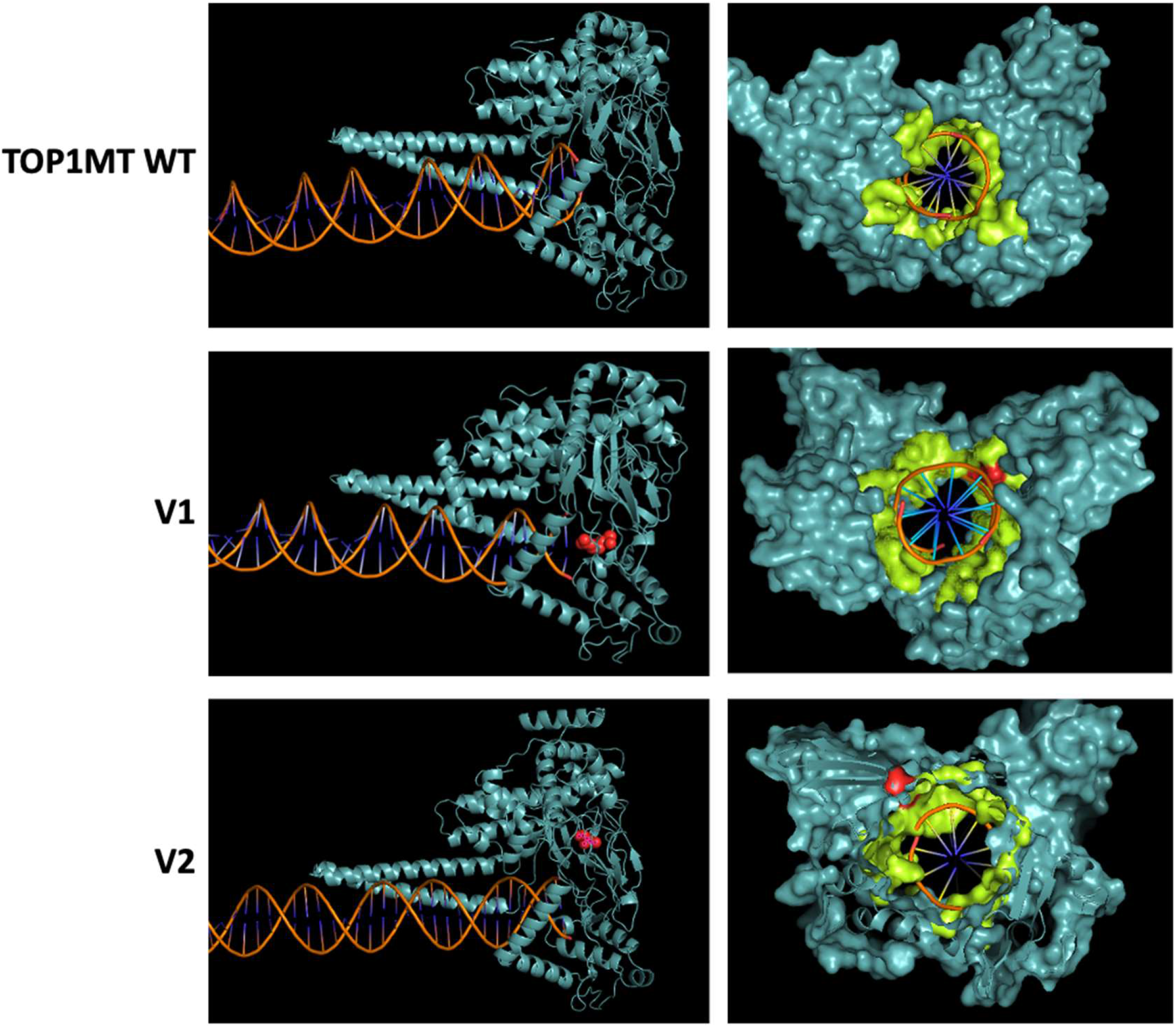
Structural models of dsDNA binding to TOP1MT variants. The structural Comparison of TOP1MT WT protein structure with its mutant forms. R198C and V338L showing the 3D model of the TOP1MT protein and the corresponding location of the amino acids of interest in red relative to the DNA binding tunnel represented in lime color.

## Discussion

To investigate whether the compound heterozygous TOP1MT variants that we identified in a newborn patient with hypertrophic cardiomyopathy could impact protein function, we examined their ability to rescue several parameters in cells lacking TOP1MT. Our results reveal several significant differences between the rescue capabilities of both V1 and V2 TOP1MT variants compared to WT TOP1MT (summarized in Table 2). The fact that V1 and V2 variants could provide partial rescue in some conditions when re-expressed indicates that they retain some functionality. However, it should be noted that the variants were overexpressed ∼10 fold above endogenous protein expression. Thus, any rescue is likely an overestimate of what would be expected at endogenous expression levels. Nonetheless, our findings clearly show that the V1 and V2 variants have impaired functionality. Intriguingly, our molecular characterization of the V1 and V2 variants also begins to provide novel mechanistic insight into the roles that TOP1MT plays in mediating mitochondrial functions. As detailed below, we noted that the V1 and V2 variants differed from each other with respect to their partial rescue for several of the parameters we investigated. These discrepancies suggest the V1 and V2 variants have distinct effects on the various TOP1MT functions.

**Table 2.** Summary of mitochondrial phenotypes analyzed. Various mitochondrial phenotypes investigated in this study, where +++, ++, +, and – represents full rescue, moderate rescue, weak rescue, or no rescue, respectively.

|  | KO-ctrl | Rescue | V1 | V2 |
| --- | --- | --- | --- | --- |
| Nucleoid size | Smaller | +++ | ++ | + |
| Supercoiling | Higher | +++ | +++ | ++ |
| mtDNA Copy number | Higher | +++ | - | +++ |
| mtDNA Repletion | Slower | +++ | - | ++ |
| Mitochondrial Transcripts | Lower | +++ | +++ | ++ |
| OXPHOS Proteins | Lower | +++ | - | - |
| Mitochondrial Oxygen consumption | Lower | +++ | + | - |
| Mitochondrial morphology | Fused | +++ | ++ | + |

### Nucleoid Size

We found that loss of TOP1MT leads to smaller nucleoids. As each mtDNA nucleoid is estimated to contain ∼1.4 copies of the mitochondrial genome (reflecting the ongoing replication of mtDNA molecules)[40], smaller nucleoid sizes likely do not reflect fewer copies of the mtDNA in each nucleoid, especially as we saw an increase in mtDNA copy number. We also showed that there were no deletions in the mtDNA, excluding another alternative explanation for smaller nucleoids. The finding of smaller nucleoids is consistent with previous work showing that loss of TOP1MT in MEF cells and murine tissues leads to increased mtDNA negative supercoiling[25], which we also observed in HCT cells lacking TOP1MT. Thus, we conclude that the smaller nucleoid sizes in TOP1MT KO cells reflect higher levels of negative supercoiling leading to more compact nucleoid structures. Both V1 and V2 variants were able to partially rescue the nucleoid size phenotype, with V1 having slightly stronger rescue, though neither was as efficient as the WT TOP1MT. In addition, both variants had a similar rescue pattern with respect to mtDNA supercoiling, with V1 showing better rescue. Collectively, these findings suggest that both V1 and V2 variants have reduced topoisomerase activity.

### mtDNA Copy Number and Replication

We also observed relevant differences in mtDNA copy number and replication.

Consistent with a trend for increased numbers of nucleoids measured using confocal microscopy, an increase in mtDNA copy number in TOP1MT KO cells was quantified by PCR. Nonetheless, despite an increase in steady state mtDNA copy number, we found that TOP1MT KO cells had decreased rates of mtDNA replication during repletion following ethidium bromide treatment, as noted in previous work[22, 36]. Curiously, while the V1 variant showed no rescue at all in the context of mtDNA copy number or replication, the V2 variant was able to restore steady state copy number and rescue mtDNA replication during repletion, though not to the same level as WT TOP1MT.

With respect to how TOP1MT impacts mtDNA copy number and replication, it is worth noting that although the human mtDNA has been studied for over four decades, there are still several unknowns about how it is regulated. Relevant to the current work, there is evidence that the mechanism of mtDNA replication varies depending on the specific conditions such as ‘maintenance replication’ to preserve copy number, compared to ‘induced replication’ to restore copy number following depletion[41]. Specifically, ‘induced replication’ appears to occur by allowing further extension of the H-strand beyond the replication termination site that usually leads to D-loop formation[42]. Notably, decreased H-strand termination also occurs under physiological conditions such as T-lymphocytes undergoing rapid cell proliferation[43].

Critically, there is a cluster of TOP1MT binding sites in the D-loop region near the site where replication termination occurs[12]. Thus, the fact that TOP1MT KO cells have decreased recovery rates under ‘induced’ conditions suggests that TOP1MT may be required to avoid H-strand termination. Conversely, TOP1MT may also inhibit ‘maintenance replication’, given the increased steady-state levels observed in TOP1MT KO cells. Combined with the ability of both V1 and V2 variants to partially rescue nucleoid size and mtDNA supercoiling, the discrepancy in the ability of V1 and V2 to rescue mtDNA copy number and repletion suggests that the ability of TOP1MT to regulate these parameters may be independent of its topoisomerase activity.

Alternatively, there could be a threshold effect with respect to the amount of supercoiling linked to mtDNA copy number and repletion, with the V1 and V2 variants on either side of the threshold.

It is also interesting that while overexpression of WT TOP1MT significantly increased mtDNA repletion rates above endogenous protein levels, it did not have a major impact on steady state mtDNA copy number, which was comparable with the endogenous protein levels. Overall, these observations suggest that TOP1MT may play distinct roles in regulating mtDNA replication and maintaining copy number. Moreover, the differences between the V1 and V2 variants could be useful in further elucidating how TOP1MT impacts mtDNA replication.

### Transcription

Another interesting finding was the fact that despite maintaining a higher steady-state mtDNA copy number, TOP1MT KO cells have reduced expression of mitochondrial-encoded mRNA genes. With respect to rescue, the V1 variant was able to restore transcript levels to a similar degree as the WT protein, while the V2 variant only had partial rescue capabilities. The similar rescue trends with respect to transcription and nucleoid size, with V1 providing better rescue than V2, suggest that TOP1MT affects transcription via its topoisomerase activity. This finding is consistent with previous reports that DNA topology impacts mitochondrial transcription *in vitro*[44-48], and that TOP1MT appears to regulate transcription via its supercoiling activity[14, 37]. Notably, our findings also suggest that a partial rescue in supercoiling activity is sufficient to rescue transcription.

Our findings that TOP1MT KO cells had reduced transcript levels contrast with previous reports that TOP1MT inhibits transcription and that KO cells had elevated transcript levels[14, 37]. The reasons for this discrepancy are likely due to differences in growth conditions, as many different parameters can impact mitochondrial function[49]. Our data suggest that different seeding densities are critical for determining how TOP1MT impacts transcript levels. However, unfortunately the seeding density was not reported for any of the previous studies. Future work will need to look at how the other roles of TOP1MT are impacted by cell density. Indeed, cell density could impact growth rate, which could be relevant to the different modes of mtDNA replication and D-loop formation discussed above, as well as the importance of TOP1MT in liver regeneration[22]. Regardless of the specific growth conditions, it is evident that TOP1MT clearly affects mitochondrial transcript levels, likely via changes in supercoiling.

### Mitochondrial Proteins and Respiration

We observed decreased protein levels of mtDNA-encoded OxPhos proteins in TOP1MT KO cells. A similar trend in reduced expression of nuclear-encoded OxPhos proteins was also observed, and possibly reflects the fact that expression of the mitochondrial-encoded subunits is rate limiting. The overall reduction in OxPhos proteins most likely explains the reduced oxygen consumption and mitochondrial impairments in TOP1MT KO cells. Although the reduced transcript levels that we observed in TOP1MT KO cells could explain the reduced translation of mtDNA-encoded genes, this conclusion is complicated by the fact that TOP1MT also has a distinct role in mediating mitochondrial translation by directly interacting with the ribosome[15].

In this regard, the differences in partial rescue of different TOP1MT functions by V1 and V2 provides unique insight, as neither V1 or V2 re-expression was sufficient to restore translation or oxygen consumption. However, given that V1 and V2 were able to restore transcript levels fully and partially, respectively, but neither fully rescued protein levels, we suggest that impaired translation in the V1 and V2 lines is largely responsible for the decreased mitochondrial function in these cells. Though little is known about exactly how TOP1MT regulates translation, our findings suggest that the role TOP1MT plays in translation is independent of its topoisomerase activity and changes in transcripts. Moreover, it is likely that the role TOP1MT plays in translation is also distinct from its roles in meditating mtDNA replication. Finally, the V1 and V2 variants, which both have deficient translation capabilities, could prove valuable tools in future studies looking at how TOP1MT regulates mitochondrial translation.

### Mitochondrial Morphology

We observed and quantified an increase in mitochondrial hyperfusion in TOP1MT KO cells, consistent with previous reports in TOP1MT knockout mouse embryonic fibroblasts[50]. Although fragmentation of mitochondrial networks is typically associated with mitochondrial dysfunction, hyperfusion of the mitochondrial network can also occur in response to various stresses[51]. While overexpression of wild-type TOP1MT restored mitochondrial morphology, V1 and V2 variants were capable of partially reverting the mitochondrial morphology. This rescue pattern resembles what we observed with respect to changes in mitochondrial nucleoid size, suggesting that changes in mtDNA nucleoid supercoiling can promote mitochondrial hyperfusion. Although the mechanism through which changes in mtDNA might impact the mitochondrial network is unknown, it is notable that a similar hyperfusion of the mitochondrial network occurs in cells with heterozygous loss of TFAM, a key mtDNA packaging protein and transcription factor[52].

### TOP1MT structural modeling

We used RoseTTAFold structure prediction to model TOP1MT binding to mtDNA. Notably, both the V1 and V2 amino acid variants localize within a core barrel that is predicted to bind dsDNA with a very strong binding affinity. However, neither amino acid substitution affected the predicted DNA binding capacity of the protein compared to the WT TOP1MT. The fact that the variants do not appear to impair DNA binding is consistent with the fact that they maintain supercoiling activity. However, given that the variants failed to rescue the translation phenotype, it seems likely that the barrel domain is important for the translational function of TOP1MT, which is a somewhat unexpected finding that begins to shed light on how TOP1MT may regulate translation. One speculative possibility is that the barrel domain of TOP1MT interacts with a mitochondrial RNA species that is import for mitochondrial translation (i.e. tRNA, rRNA, or mRNA). In this regard, RNA species can for double-stranded secondary structures that can resemble DNA, while other topoisomerase proteins can bind RNA[53-56].

### TOP1MT in Human Disease

Although the *TOP1MT* gene is not formally recognized as a human disease gene, our novel findings, combined with previous work by others, strengthen the argument that TOP1MT should be considered for its role in human disease. In a study of 40 patients with disorders of mtDNA maintenance, 39 candidate genes linked to mtDNA maintenance were sequenced, and two predicted pathogenic heterozygous variants in TOP1MT were identified[57]. However, given the limited number of genes sequenced at the time, and a lack of mechanistic studies, a direct correlation between the patient phenotype and pathogenicity of the candidate variants was not established. In a separate study, 30 potentially deleterious TOP1MT variants occurring in the normal population were reported[24], including the V2 variant that we studied here. Again, however, no functional characterization of these variants was performed. Meanwhile, a functional study of TOP1MT single nucleotide polymorphisms (SNPs), found that some SNPs have reduced DNA relaxation activities *in vitro*[58]. Notably, these SNPs include a rare variant (E168G) found in a patient with mitochondrial deficiency syndrome, and two common single nucleotide variants (V256I and R525W) that are most often mutually exclusive, suggesting there is selection against both variants being present together.

Mouse studies also support a role for TOP1MT in maintaining mitochondrial function. Although *Top1mt* knockout mice are viable and fertile[13], they are sensitive to doxorubicin-induced cardiotoxicity[24], have impaired liver regeneration[22], and embryonic fibroblasts from these mice have defective mitochondrial function[50]. Our findings are certainly consistent with the notion that the V1 and V2 variants contribute to mitochondrial dysfunction, most likely through some combination of impaired topoisomerase activity, altered mtDNA replication, and reduced mitochondrial translation. However, we cannot definitively ascribe the cardiomyopathy in our patient directly to mitochondrial dysfunction due to the V1 and V2. Nonetheless, the increased cardiotoxicity of *Top1mt* KO mice to doxorubicin emphasizes the importance of TOP1MT to cardiac function. Additionally, mtDNA genomes in cardiac tissue are organized into complex catenated networks[59], which may make them more dependent on topoisomerase activity. Meanwhile, there is a significant transition in cardiac function at birth that has critical implications for mitochondrial function, as the heart adapts to changes in oxygen levels, increased energy demand, and a metabolic shift from glycolysis to beta-oxidation. Thus, we suggest that impaired TOP1MT activity, combined with the dependence of the heart on mitochondrial function, is likely to be contributing to the cardiomyopathy in the patient.

## Conclusions

Overall, our characterization of the V1 and V2 variants support the argument that TOP1MT should be considered as a mitochondrial disease gene. Additionally, characterization of our stable cells lines provides insight into the multiple distinct and interrelated functions of TOP1MT including relaxing mtDNA supercoiling, regulating mtDNA transcript levels, mediating mitochondrial replication, and promoting mitochondrial protein translation. By comparing the distinct rescue patterns of the V1 and V2 variants, our findings support that idea that the topoisomerase activity of TOP1MT is related to its role in regulating mitochondrial transcription, as well as changes in mitochondrial morphology. Meanwhile, the roles that TOP1MT plays in mtDNA replication, maintenance of mtDNA copy number, and mitochondrial translation appear to be separate from each other, as well as independent of the topoisomerase activity. Future characterization of the V1 and V2 variants, as well as other TOP1MT variants, will likely contribute to delineating the roles that TOP1MT plays in maintaining mitochondrial function and its role in human disease.

## Data Availability

The data that support the findings of this work are available from the corresponding author upon request.

## Author contributions

I.A.K. conceived the study, designed, performed and analyzed experiments and wrote the manuscript. J.D performed experiments. S.S. generated the structural models. M.K. identified the *TOP1MT* patient variants. H.Z., S.H. and Y.P. generated the HCT116 TOP1MT KO cells, shared the TOP1MT cDNA, and provided input on the project. A.K. performed the clinical workup of the patient, provided input on the project, and edited the manuscript. T.S. conceived and designed the study, analyzed experiments, supervised the study, and wrote the manuscript.

