## Supplemental Data for "Functional characterization of two variants in the mitochondrial topoisomerase gene TOP1MT that impact regulation of the mitochondrial genome"

### S1. Characterization of HCT116 control and TOP1MT-KO cells.

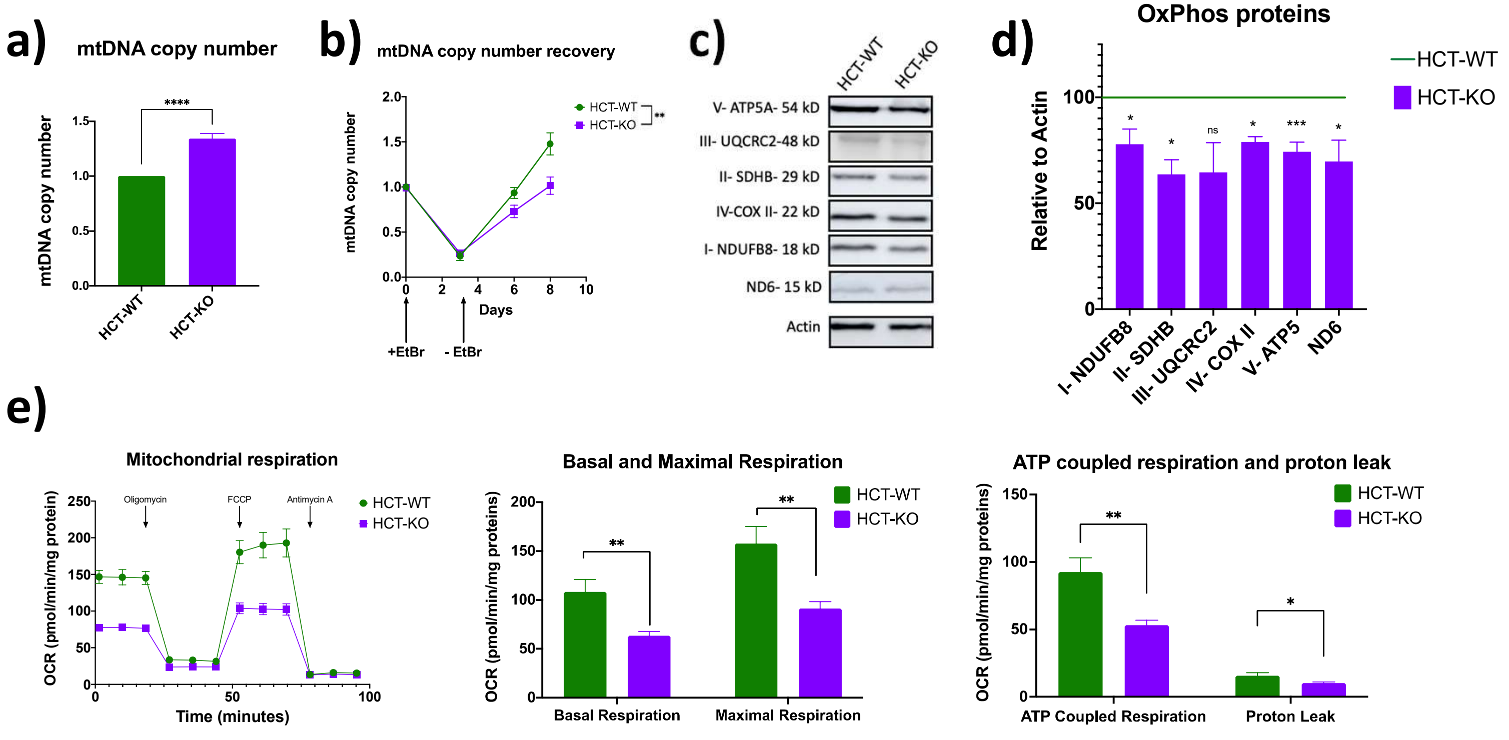


**S1. Characterization of HCT116 control and TOP1MT-KO cells**. **a)** Relative mtDNA copy number, determined by qPCR relative to 18S, is higher in HCT-KO cells compared to HCT-WT cells. **b)** mtDNA copy number increases more slowly in HCT-KO cells compared to HCT-WT cells following EtBr depletion for 3 days. **c)** Representative western blots shows reduced levels of the indicated OXPHOS complex proteins. **d)** Quantification of western blots of OXPHOS proteins as in (b), from three independent experiments, corrected to Actin as a load control. Reduction in OXPHOS proteins encoded by both nuclear (i.e. NDUFB8, SDHB, UQCRC2, ATP5) and mtDNA (i.e. COXII, ND6) genomes was observed in cells lacking TOP1MT. **e)** Mitochondrial respiration analysis shows lower oxygen consumption rates in KO cells at both basal and maximal respirations which is reflected on lower ATP coupled respiration and decreased proton leak. All statistical analysis were done using unpaired student t-test and p values * <0.05, ** <0.01, *** <0.001 and **** <0.0001. 'ns' signifies no significant differences between indicated groups. Error bars represent standard error of mean.

### S2. Nucleoid sizes in TOP1MT cell lines using ant-DNA antibody.

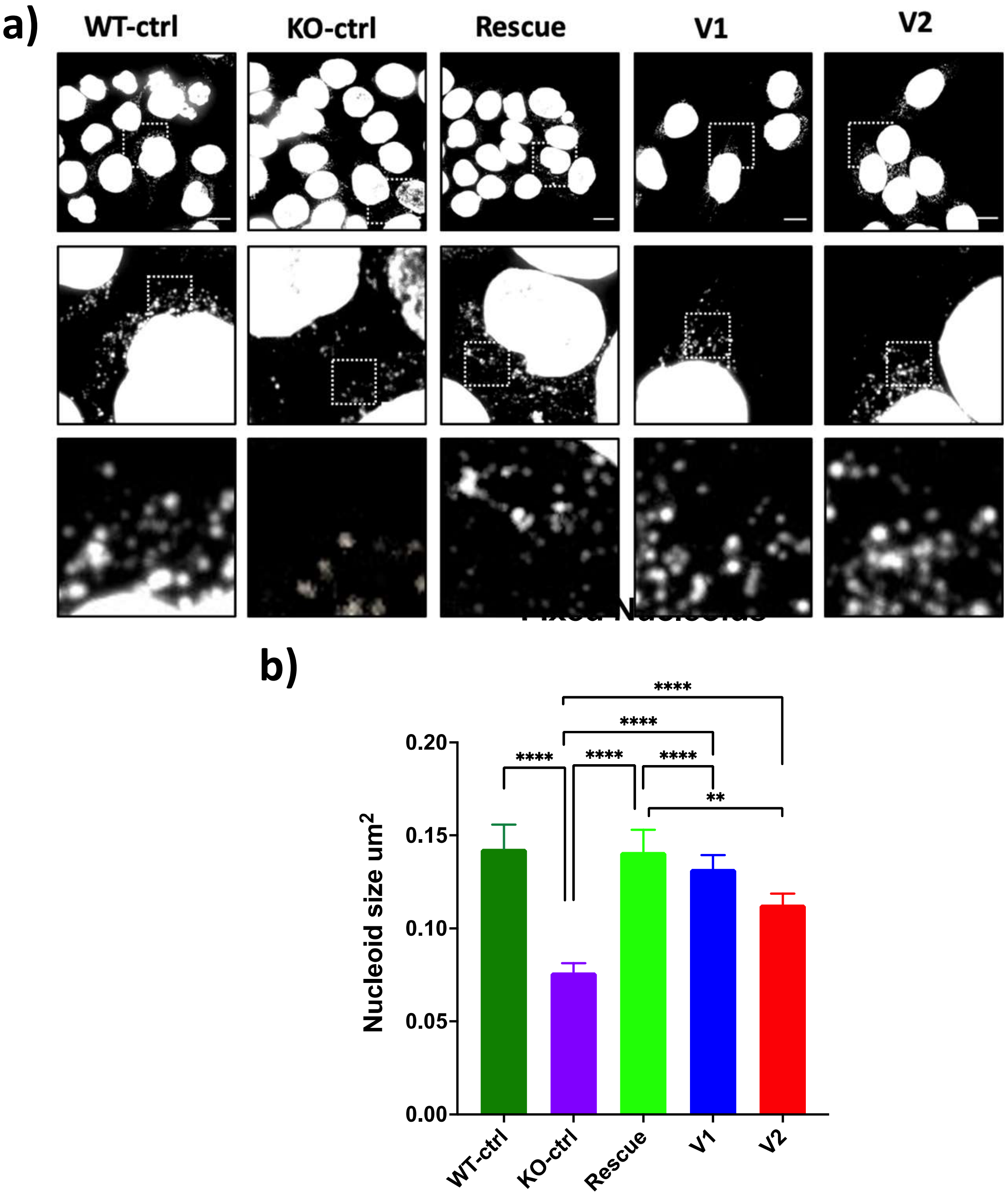


**S2. Nucleoid sizes in TOP1MT cell lines using ant-DNA antibody. a)** Quantification of mtDNA nucleoid sizes in cells fixed and labeled with anti-DNA antibody. Data represents combined nucleoid sizes from at least 15 cells for each group. Error bars represent standard error of mean. P-values were determined by a Kolmogorov-Smirnov test for all measured nucleoids, with ** <0.01 and **** <0.0001.

### S3. Expression of mtDNA encoded genes in different seeding densities of KO cells relative to WT cells

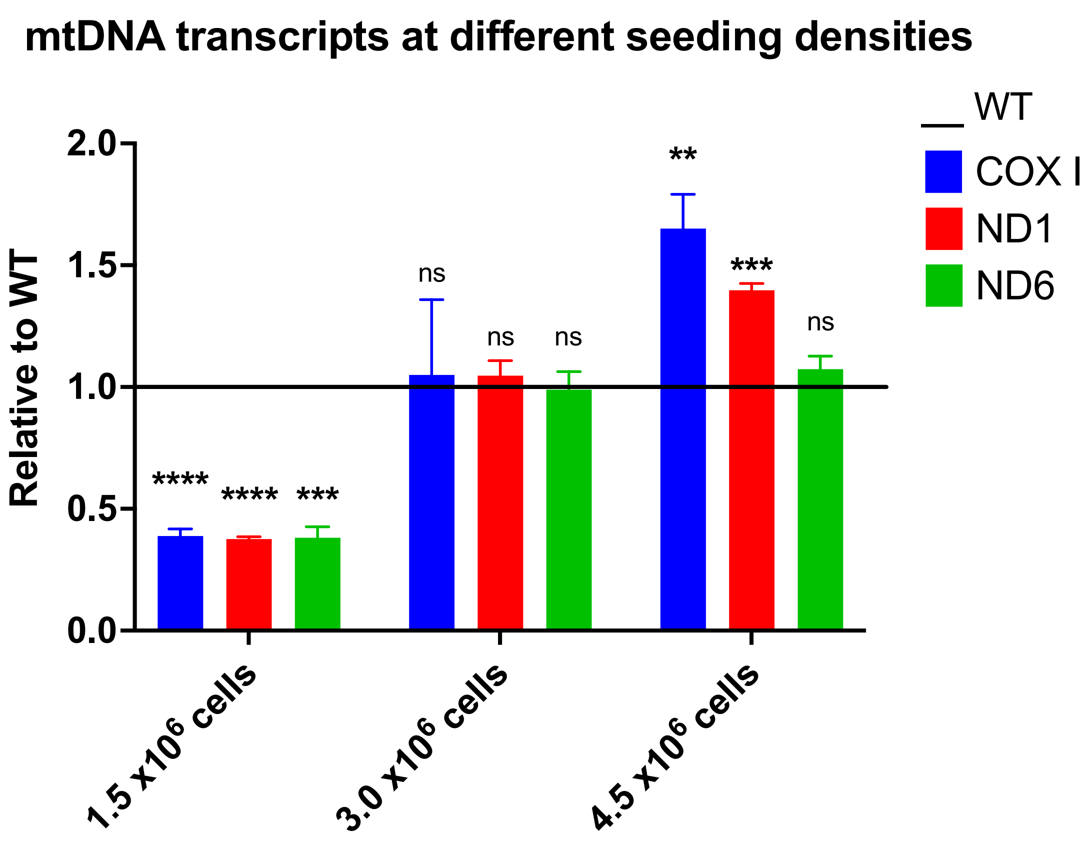


**S3. Expression of mtDNA encoded genes in different seeding densities of KO cells relative to WT cells a)** Relative expression of mtDNA transcripts from KO cells seeded at 1.5 x $10^6$, 3 x $10^6$ and 4.5 x $10^6$ cells compared to WT cells seeded at the same density. Data represented as the transcript fold change in KO relative to WT cells from three biological replicates. Error bars represent standard error of mean. P-values were determined using unpaired student t-test and p values ** <0.01 and *** <0.001, **** <0.0001 and 'ns' signifies no significant differences between indicated groups. Error bars represent standard error of mean.
